# A phylogenetic and transcriptomic study of the β-1,3-glucanase family in tomato identifies candidate targets for fruit improvement

**DOI:** 10.1101/2021.09.29.462359

**Authors:** Candelas Paniagua, Louisa Perry, Yoselin Benitez-Alfonso

## Abstract

Tomato, *Solanum lycopersicum*, is one of the most cultivated fruits. However, between one-quarter and half of their production is lost during transport and storage. Modifications in cell walls, and specifically pectin composition, delay fruit softening but, so far, the impact of callose metabolism in this process has not been investigated. Callose accumulates in cell walls around plasmodesmata to modify symplasmic transport. It also plays a role in reinforcing cell walls in response to bruising or pathogen invasion. The aim of this work is to identify cell wall β-1,3-glucanases expressed in tomato fruit that can be used as targets to modify callose accumulation during ripening. A phylogenetic analysis identified fifty candidate β -1,3-glucanases in tomato distributed in three clusters (α, β and γ) with evolutionary relations previously characterised in the model *Arabidopsis thaliana*. Analysis of tomato microarray data indicates different regulatory patterns: the expression of a subset of enzymes in cluster α decreased during ripening, while enzymes in cluster β and γ displayed higher expression in white-red stages. qRT-PCR experiments confirm the differential regulation of enzymes in different clusters suggesting evolutionary divergences that correlate with differences in their predicted localization and function. The potential to exploit this information in the selection of targets to modify cell walls and fruit development is discussed.

## 1. Introduction

The tomato fruit, *Solanum lycopersicum*, is, according to the Food and Agriculture Organization (FAO), the second most cultivated fruit worldwide, with 243 million tonnes of fruit produced in 2018 [1]. Exports of tomato fruits from EU countries accounted for 17.7% of all vegetable outputs in the period 2015–2017 [2]. In spite of this, losses in production are considerable and are estimated to range between 24 and 50% in a single year [3]. Poor conditions during transport and storage from field to market are the main causes for these losses, which are influenced by a lack of temperature control, physical injury, pests and disease [3]. Understanding the biological components and processes that affect the shelf-life of the tomato fruit is essential to design new strategies to reduce these losses.

One of the processes that influences the shelf-life of tomato fruit is their rapid softening. Softening is a biological process that involves the disassembly of the cell wall (CW), changes in cell turgor, fruit–water status, hydrostatic pressure and in the accumulation and distribution of osmotically active solutes [4]. Delaying this process is one of the major targets in fruit breeding programmes. Research on this area focuses on introducing changes in the expression of specific cell wall-modifying enzymes (CWME) that target major cell wall components (such as pectins and cellulose) [5]. However, the role of minor components, such as callose (β-1,3-glucan), in tomato ripening and shelf-life remains unclear.

Callose is a plant cell wall polysaccharide synthesised by callose synthases (Cals) and hydrolysed by β-1,3-glucanases (BG) [6,7]. Callose synthesis/degradation is involved in disease response, in the control of intercellular transport and in the regulation of cell wall integrity and mechanical properties [8]. The objective of this study was to identify candidate BG participating in tomato ripening. We analysed the glycosyl hydrolase family 17 (GH17) which comprises all BG proteins identified so far. Structural features (presence of protein sequence domains and catalytic residues) were predicted and phylogenetic comparison carried out to screen for candidate BGs in tomato. The expression of the genes in tomato was studied using publicly available transcriptomic data and experimental determinations. The results reveal differential regulatory patterns for enzymes grouped in distinct clusters, likely associated with enzymatic function in different cellular micro-domains or in response to different environmental cues. The potential to exploit these differences in the selection of targets for tomato fruit improvement is discussed.

## 2. Methods

### 2.1. Isolation of β-1,3-Glucanase Gene Sequences and Prediction of Domains and Catalytic Residues

Glycoside hydrolase family 17 (GH17) proteins were identified in *Arabidopsis thaliana* and *Solanum lycopersicum* using Phytozome [9] and the search term ‘glyco_hydro_17’. Presence of a GH17 domain was confirmed in these sequences using the SMART [10], Conserved domain [11] and Interpro [12] prediction tools. To identify and remove redundant sequences, the UGENE [13] bioinformatics software was used together with the MUSCLE [14] alignment function.

Sequence features previously associated with the BG family, such a signal peptide (SP), glycosylphosphatidylinositol-anchors (GPI-anchors) and X8 (CBM43) domains were identified using prediction platforms [15]. Presence of signal peptides (SP) was predicted using Aramemnon [16], SignalP v5.0 [17], Spoctopus [18], Predisi [19], SMART [10] and Interpro [12]. Presence of GPI anchor was predicted using KohGPI [20], PredGPI [21] and BigP [22]. Presence of X8 domains was predicted using SMART [23], NCBI CDS [11] and Interpro [12]. Presence of these sequence signatures were considered positive when predicted by at least one of these databases.

The catalytic domain of GH17 family of proteins display two catalytic glutamate (E) residues which act as a proton donor (E94) and nucleophile (E236), important to cleave substrates that contain a beta-1,3-linkage [24]. The presence and position of these catalytic residues was predicted as described before [15].

### 2.2. Comparative Phylogenetic Analysis of Tomato and Arabidopsis genes

*Arabidopsis thaliana* and *Solanum lycopersicum* GH17 were aligned using MUSCLE [14]. Phylogenetic analysis was carried out in MEGA-X 10.1.840 software using the Jones–Taylor–Thornton (JTT) model. Conditions and restrictions were varied to produce a range of dendrograms with altered bootstrap values (between 100 and 1000), branch swap filter settings (none or very weak) and ML Heuristic Method settings (Nearest neighbour interchange (NNI)) or extensive Subtree pruning re-grafting (SPR level 5). *Arabidopsis thaliana* GH17 genes with common names were annotated. Confidence and clustering was based on comparison with Doxey et al. (2007) [25].

### 2.3. In silico Gene Expression Analysis

Tissue and organ expression data for tomato GH17 genes was obtained from Genevestigator [26] and compared to the ePlant BAR [27]database. Expression in pericarp at different stages of fruit development was obtained from the SGN Tomato Expression Atlas (TEA) [28] and TomExpress [29]. Protein intracellular localisations were predicted using PSORT [30] and WoLF [30] prediction tools.

### 2.4. RNA isolation and Expression analysis by quantitative real-time PCR

Total RNA was isolated from pools of tomato fruit (var. MicroTom) collected at different stages (from 3 days post-anthesis to mature red fruit). RNA was isolated and purified using the total RNA extraction and purification kit (Monarch, NEB). RNA concentration and integrity were calculated using a Nanodrop and agarose gel electrophoresis. RNA sample was treated with DNase I to remove genomic DNA and retro transcribed into cDNA using Quantitect Reverse Transcription Kit (Qiagen).

Gene expression analysis of selected genes was performed using Quantitative real-time iCycler (BioRad) and QuantiTect SYBR® Green PCR Kits (Qiagen). Gene-specific primers were designed using Primer3 and blast was carried out using NCBI BLAST to determine any miss-priming (Table S1). When possible, primers were designed to flank introns aiming to determine and avoid the presence of gDNA amplification. Each reaction was performed in triplicate and Actin (ACT, Solyc11g005330) and the Clathrin adaptor complex medium subunit (CAC, Solyc08g006960) were used as reference genes [31]. Amplification efficiencies were near 100% as calculated using the Livak method [32].

## 3. Results

### 3.1. Phylogenetic Relations identify Tomato orthologues to Arabidopsis BG

Gene sequences encoding members of the GH17 family in tomato were identified using the Phytozome database [9]. 51 genes were identified but two: Solyc01g059980 and Solyc01g060020, encoded the same protein sequence. BG enzymes that regulate plasmodesmata or the response to pathogen invasion have been identified in Arabidopsis [8,33–35]. To identify candidate tomato orthologues, a phylogenetic tree was obtained as described in Material and Methods (Figure 1). Three clusters—α, β and γ— were identified to group the tomato genes as described previously for Arabidopsis [25]. Characterized Arabidopsis BGs are annotated in the tree. These comprise the plasmodesmata proteins PdBG1, PdBG2, PdBG3 and ATBG_PAP in cluster α, the signalling proteins ZET and ZETH in cluster β and the defense proteins AtBG1-BG5 in cluster γ [8,33–36].

**Figure 1.**
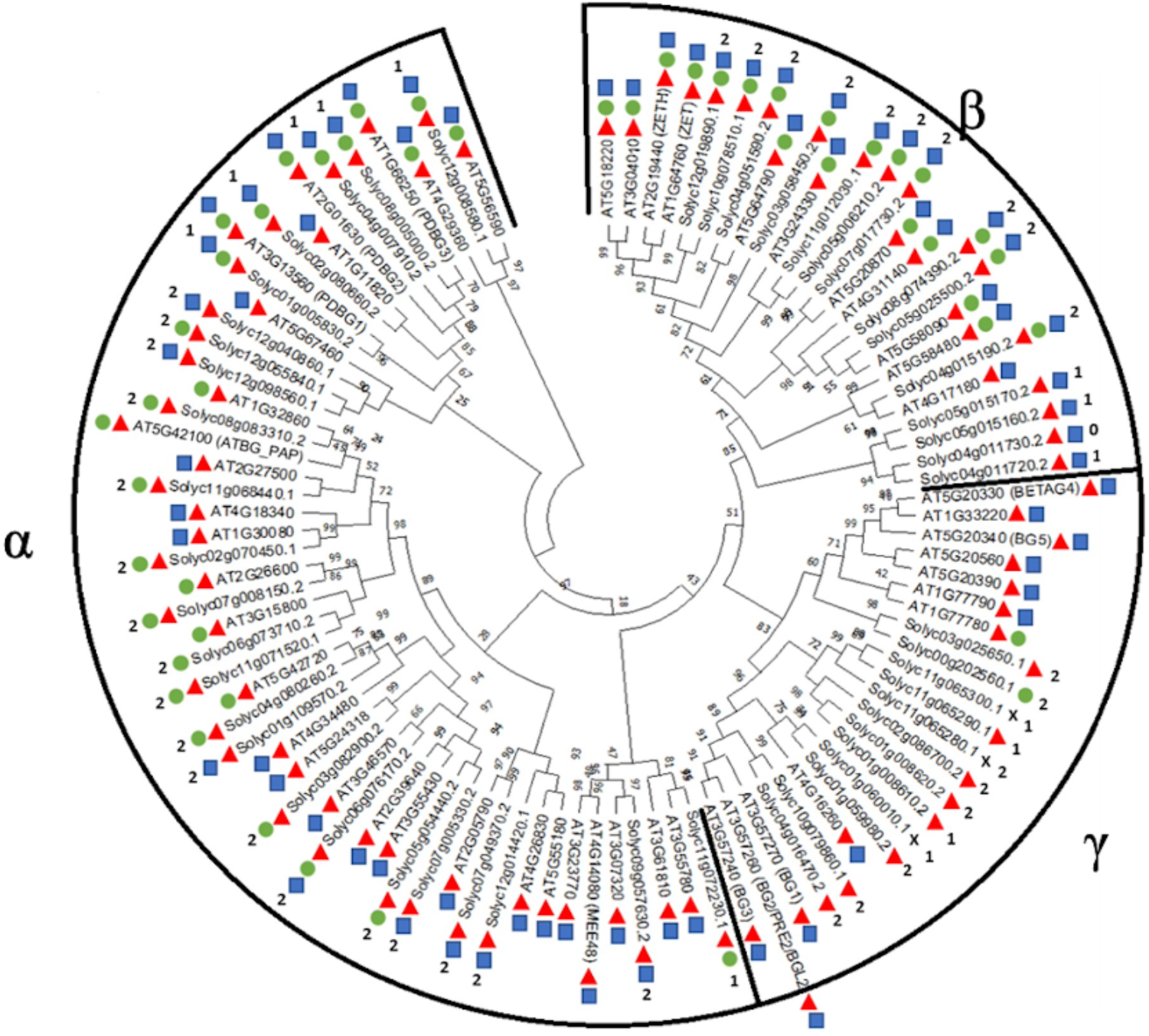
Phylogenetic relations and structural features of the glycosyl hydrolase family 17 in tomato and Arabidopsis. Amino acid sequences of predicted GH17 genes were aligned and a phylogenetic tree was created using MEGA-X40 (Jones-Taylor-Thornton model, bootstrap value of 1000). Posterior probability is displayed in the tree branches. Three clades (α, β and γ) are defined as described before in Arabidopsis. Predicted domains for each gene are indicated by the coloured spots: red triangle = signal peptide, green circle = GPI-anchor, blue square = X8 domain, ‘x’ = no domain is predicted. The number of glutamate residues in the catalytic domains (1 or 2) are also indicated.

The presence of signal peptide (SP), GPI anchor, X8 (carbohydrate binding) domain and of conserve glutamate (E) residues in the catalytic domain are features of previously characterized BG [25]. Predictions of these features, using online platforms, classified the tomato protein sequences in types 0-5 (Supplemental table 2). Results from these predictions were used to annotate the tree (Figure 1).

Most tomato genes (45 out of 50) encoded proteins with a predicted SP (by at least one of the six platforms used), suggesting these are mainly secreted proteins (Supplemental table 2). The presence of GPI anchor (which targets proteins to the membrane) was predicted for 28 out of the 50 tomato GH17 proteins. Lack of GPI domain in proteins with SP (type 4) suggests proteins likely secreted to the apoplast. With only one exception, all cluster γ proteins lacked predicted X8 binding domain whereas all in the β cluster were positive for this prediction. Cluster α was the most extensive and varied (in terms of structural predictions) but only one protein within this cluster lacked a SP prediction. Proteins in cluster α have either a GPI or X8 domain or both.

Research identified in GH17 proteins two glutamate residues that characterise the catalytic domain of glycosyl hydrolases [37]. Searching the presence of these residues in the catalytic domain reveals proteins that lacked one or both residues (Supplemental table 2). Of the 50 tomato sequences, 36 displayed both residues, 13 showed the presence of only one, and one lacked both aminoacids in the conserved position. The absence of one of these residues is not sufficient to classify these as ‘dead’ enzyme as characterization of potato (*Solanum tuberosum*) endo-BG with a mutation in the active site showed residual endoglucanase activity [24] .

The results indicate that predicted GH17 proteins in tomato have diverse sequence signatures, suggesting differences in localization and function. It also highlights tomato family members closely related to Arabidopsis BG proteins with previously characterized roles in plasmodesmata regulation, defense responses and signalling.

### 3.2. The Expression of GH17 Genes Vary Across Different Fruit Developmental Stages

Tissue expression data for the 50 predicted tomato GH17 genes was extracted from the Genevestigator [26] database. The results indicate that the expression of the tomato genes vary in different tissues and organs (Figure 2).

**Figure 2.**
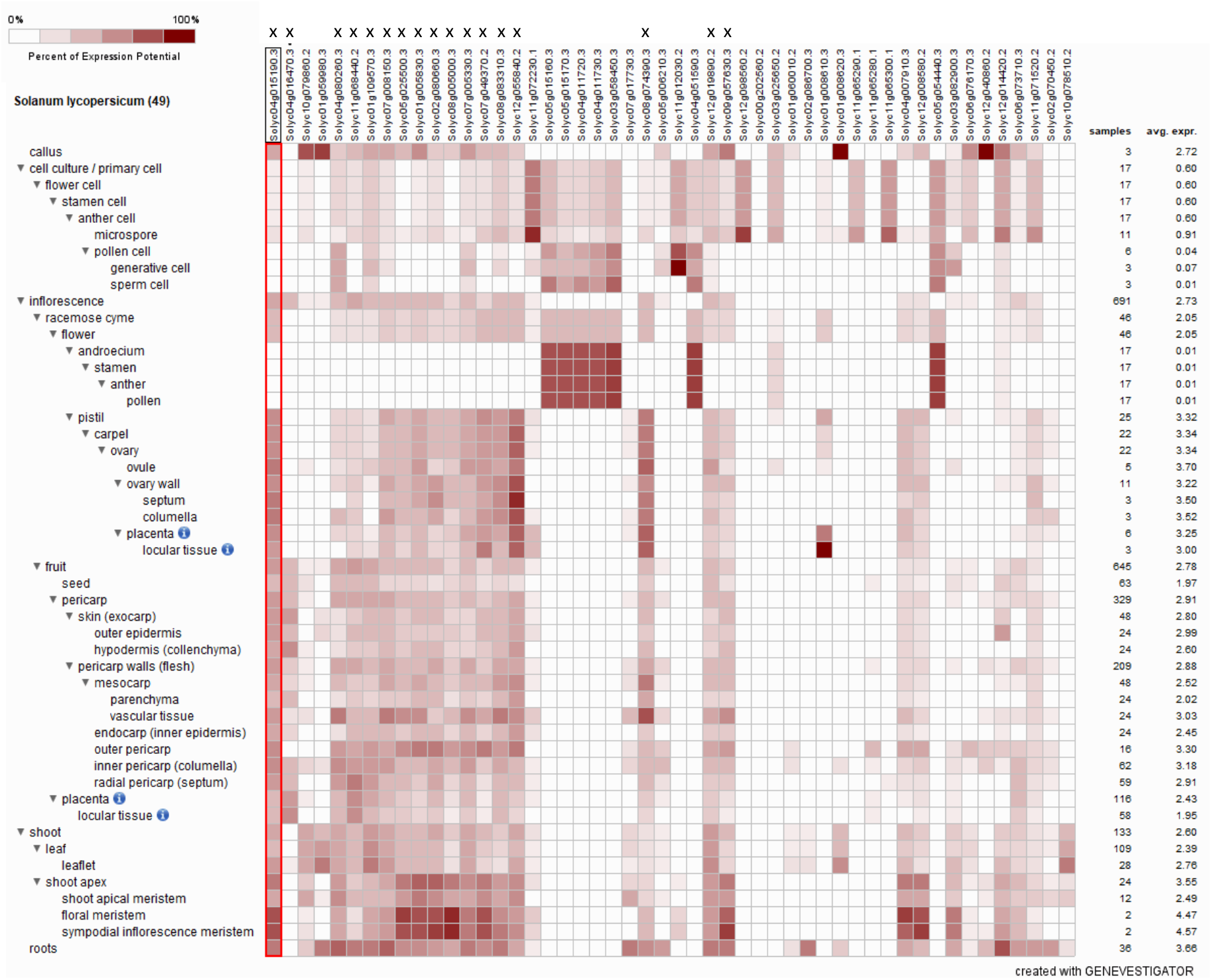
Plant tissue expression data for GH17 genes identified in tomato. Gene expression data in different tissues (49 anatomical parts) including primary cell, inflorescence, fruit, shoot, leaf, and root tissues was extracted from the Genevestigator database. The colour gradient (white to dark maroon) represents expression level low to high (from data selection: SL_mRNASeq_Tomato_GL-0). Genes expressed in fruits are labelled with an (x) on the top row.

We selected those genes with significant expression in fruits in relation to other organs. 17 proteins were identified with medium-high expression in tomato fruit (Fig. 2 and Table S2). Solyc04g016470 is the only gene expressed specifically in tomato fruits and no other organs. To identify the patterns of expression across the fruit stages, microarray data was obtained from the TEA database and the results represented in a heatmap using Clustvis [38] (Figure 3).

**Figure 3.**
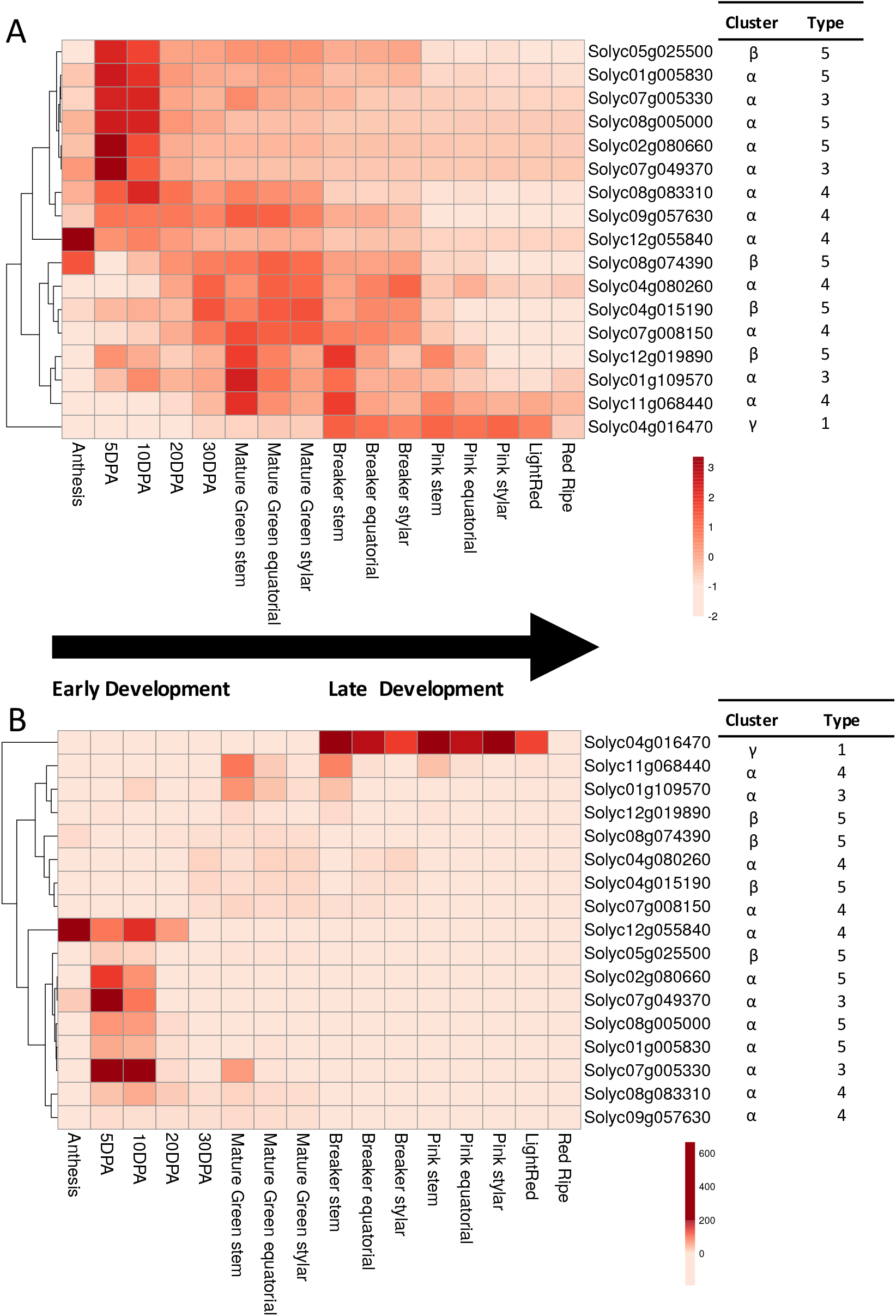
Gene expression analysis of GH17 genes in tomato pericarp during fruit development. The heatmaps show relative expression (red=high, white =low) of GH17 genes in fruit pericarp from anthesis to red ripe (DPA = days post-anthesis) based on data extracted from the SGN-TEA database. The heatmap was performed using Clustvis web tool either applying unit variance scaling to row data (**A**) or representing data with no scaling (**B**). See Table S3 for raw values. Rows are clustered using correlation distance and average linkage. The cluster and type (as described in table S2) is indicated on the right of the sequence ID.

Two main patterns of expression are identified: a subset of genes (mostly from cluster α) shows high expression early in fruit development whereas the expression of a second subset (mostly from cluster β and γ) is low at the start but increases as the fruit matures (Fig. 3A)

When analysing raw expression data (not scaling), major differences are noted for gene Solyc04g016470 (belonging to cluster γ), which is no expressed in anthesis but ramps up expression to >400 RPM during the breaker stage (Fig. 3B and table S3). On the other hand, Solyc12g055840 (cluster α) is highly expressed (805.68 RPM) during anthesis but expression is as low as 4.9 RPM in pink fruits. Predictions of structural features indicate that Solyc04g016470 is a secreted protein whereas Solyc12g055840 is GPI anchored to the plasma membrane (Fig. 3). Few other proteins are moderately well expressed early in fruit development. These include cluster α genes Solyc02g080660, Solyc07g049370, Solyc08g005000, Solyc01g005830 and Solyc07g005330 (Fig. 3B). GH17 proteins in cluster β display low expression at all stages.

The results suggest different patterns of expression for BG proteins in different clusters likely associated with distinct localization and function during tomato fruit development.

### 3.3. Quantitative RT-PCR corroborates differential expression profile for GH17 Genes during Tomato Fruit Development

To corroborate the microarray results, qRT-PCR experiments were performed for a subset of tomato genes (Figure 4). Fruits of the Microtom variety were collected and total RNA extracted from eight different developmental stages: from 3 DPA to red ripe. Solyc04g016470 and Solyc12g055840 were selected for this analysis as the highest expressed genes in the SGN fruit pericarp dataset (Table S3). Four other cluster α genes were selected: Solyc01g005830, Solyc02g080660 and Solyc11g068440 which displayed moderate levels of expression and Solyc01g109570 which expression is low. By selecting this varied list of genes, the aim was to confirm the validity of the microarray data as a whole. Primers were designed to amplify the transcripts which were analysed by qRT-PCR using ACT and CAC as reference genes (table S1).

**Figure 4.**
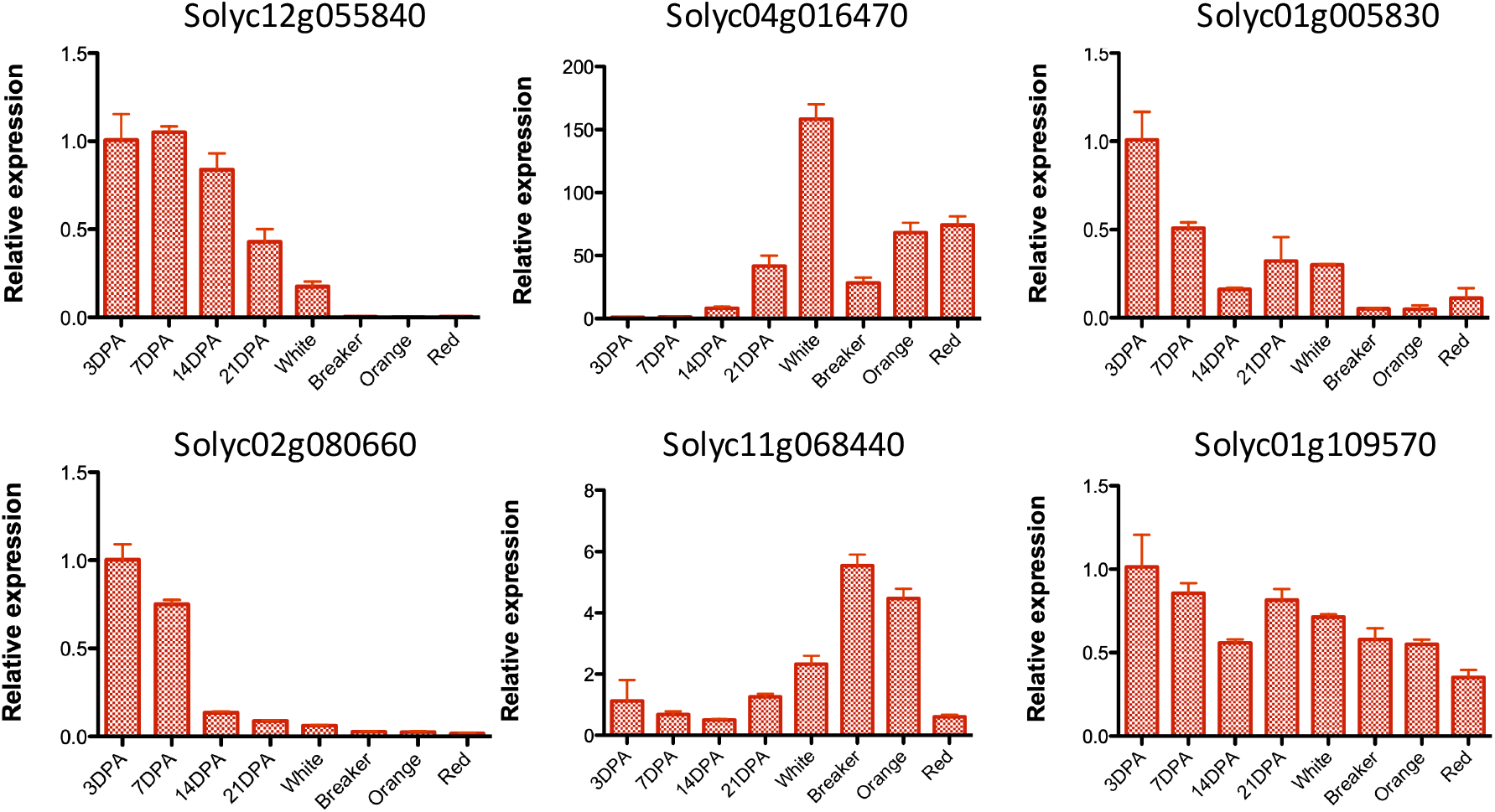
Expression analysis of a subset of GH17 genes expressed during fruit development. The relative expression of selected GH17 genes in tomato (Microtom variety) was determined using qRT-PCR and actin as reference gene. Expression values are relative to stage 3 DPA (DPA = days postanthesis). Error bars represent standard deviation of the mean for three replicas.

The results confirmed the patterns of expression observed in the microarray, Solyc04g016470 induced at white and late stages and Solyc12g055840 showing a strong decrease in expression from 14 DPA (Figure 4). Solyc02g080660 and Solyc01g005830 also showed a decrease in expression as fruit develops, corroborating the microarray results. Solyc11g068440 showed an increase expression during maturation as in the microarray data, but the increase is moderate-low when compared to Solyc04g016470 (note relative expression values). The expression of Solyc01g109570 was very low and did not change significantly across all fruit developmental stages corroborating again the expression on microarray (Supplemental table 3).

Together the results corroborate the expression detected in microarrays and indicate a different pattern of expression during tomato fruit development for GH17 proteins. In general, cluster α proteins seem to control early development whereas Solyc04g016470 grouped in cluster γ likely play a role in maturation and ripening.

## 4. Discussion

In this publication, we report the analysis of the gene family GH17 (comprising all characterized BG so far) at different stages during the development of tomato fruit. Using a combination of phylogeny and transcriptomic analysis, we found fifty GH17 enzymes in tomato, 17 of which were expressed in fruits according to microarray data. Phylogenetic analysis revealed that tomato genes are distributed in three distinct clusters (α, β and γ) which suggest both, redundancy, and evolutionary diversification of function. Previous analysis of cluster α in Arabidopsis identified proteins that localise at cell wall domains surrounding plasmodesmata, while enzymes in cluster γ included the pathogenesis-induced proteins AtBG1–BG5 [25]. Phylogenetic relations and structural analysis indicate that the tomato genes Solyc01g005830, Solyc04g007910, Solyc08g005000, Solyc02g080660 and Solyc12g055840 are GPI anchored membrane proteins likely orthologous to PDBG1, PDBG2 and PDBG3 [36] (Fig. 1). On the other hand, the tomato genes Solyc10g079860 and Solyc04g016470 are apoplastic proteins closely related to pathogenesis-related (PR) proteins AtBG1, BG2, and BG3 grouped in cluster γ [24]. Interestingly, five of the genes mentioned above were expressed in tomato fruit pericarp but with very different levels and pattern of expressions. PDBG orthologues Solyc01g005830 Solyc08g005000, Solyc02g080660 and Solyc12g055840 display the highest expression early in tomato fruit development, whereas the AtBG1-3 orthologue Solyc04g016470 was expressed at the transition to maturation stage (Fig. 3). Experimental work using qRT-PCR confirms the distinct pattern of expression for Solyc04g016470, Solyc02g080660 and Solyc12g055840 suggesting parallelisms between gene expression, protein localization and predicted function.

The expression of plasmodesmata located BG in tomato might be linked to the symplasmic unloading of sugars and signalling molecules into the ovaries and fruit tissue required to induce fruit formation and control early development. Callose restricts plasmodesmata transport thus the hydrolytic activity of these BGs might be required for symplasmic communication and phloem, post-phloem unloading [8]. An study in Chinese jujube suggests that high symplasmic communication correlates with high sugar content in the cultivated variety [39]. In tomato fruits a shift in the sugar unloading pathway (from symplasmic to apoplasmic) is described at the onset of ripening [40]. This transition might explain the reduction in the expression of these candidate plasmodesmata enzymes later in development (white and onwards) [40].

On the other hand, the ‘relatively late’ expression of pathogenesis and stress-related enzymes (grouped in cluster γ) might be linked to the start of fruit softening, leading to increased susceptibility to pathogen attack [41]. Comparative analysis in three banana cultivars: Rasthali, Kanthali and Monthan, suggests a correlation between the expression of PR BG and differences in the rate of softening [42]. The expression of cluster γ BG might also have a protective effect against fungal infection [43]. GH17 proteins isolated from different species, including alfalfa, barley, soybean, tobacco and wheat, were co-expressed with antifungal proteins (such as chitinases and peroxidases) to improve pathogen resistance in various crop species [44]. In tomato, co-expression of the tobacco BG (GLU) and alfalfa defensin gene (alfAFP) enhanced resistant to the pathogen *Ralstonia solanacearum* [45]. Digestion of beta 1,3 glucans from pathogen cell wall also serve as elicitors to activates structural defence responses [46]. Studies in tobacco leaf (*Nicotiana tabacum*) indicate that β-1,3 glucan oligomers induce the synthesis of PR proteins for the development of resistance against infection with the bacterial pathogen *Erwinia carotovora* subsp. *carotovora* [47,48]. In wheat, beta-1,3-glucan oligomers cleaved from the pathogen cell walls act as signal to prevent *Septoria tritici* colonization [49].

Based on our results, we propose that differences in the expression and localization of BG enzymes need to be considered for the selection of appropriate targets to improve tomato fruits. We have identified a group of enzymes likely controlling plasmodesmata and symplasmic unloading early in development and a separate enzyme likely involved in the defense mechanism late during ripening. Understanding the biological components and processes that affect the fruit quality, quantity and shelf-life is essential to design new strategies to improve their cultivation and availability. Current work focuses in understanding the possible role of these callose degrading enzymes in phloem unloading, cell-to-communication and defense in order to determine their suitability as targets for translational approaches.

## Author Contributions

C.P. and Y.B.-A. conceived and designed the experiments and the project; L.P. performed the experiments; L.P., C.P. and Y.B.-A. analysed the data; L.P., C.P. and Y.B.-A. wrote the paper. All authors have read and agreed to the published version of the manuscript.

## Acknowledgments

This project has received funding from the European Union’s Horizon 2020 research and innovation programme under the Marie Skłodowska-Curie grant agreement No. 839086. We would like to thank Sam Amsbury and Philip Kirk for help with transcriptomic analysis.

