## Supplemental tables for "A phylogenetic and transcriptomic study of the β-1,3-glucanase family in tomato identifies candidate targets for fruit improvement"

**Supplementary Table 1. List of primers used in qRT-PCR analysis and amplicon size.**

| Gene ID | Melting Temperature (C) | Product length (bp) | Primer Sequence (5'-3') |
|---|---|---|---|
| Solyc04g016470 Fw | 56.28 | 85 | CACTAGTTACCCCAGATTTTACAG |
| Solyc04g016470 Rv | 57.58 | | TCTGCTGGAGATGGTAATCC |
| Solyc02g080660 Fw | 60.69 | 155 | CAACCAAATTCTTGCTATTGGG |
| Solyc02g080660 Rv | 62.95 | | TCATTGCATCAAGCACATTGG |
| Solyc12g055840 Fw | 60.11 | 191 | TCGATGCGGTTCATTCTGCT |
| Solyc12g055840 Rv | 60.04 | | GCTGTTTGGCTTCAACGGAG |
| Solyc01g005830 Fw | 59.87 | 150 | ACTGTAGACTTGTGTCTTGTGCT |
| Solyc01g005830 Rv | 58.28 | | AAGTGGACAAAGCTGACAACA |
| Solyc01g109570 Fw | 57.71 | 196 | TGTCCATTTTGTCCCAGTCC |
| Solyc01g109570 Rv | 56.98 | | CCAGCATTTGGTTGGAAGAG |
| Solyc11g068440 Fw | 60.04 | 142 | AACGTGTTAACCGCTTTCGC |
| Solyc11g068440 Rv | 60.25 | | ACAAGAGCACGGTACACACC |
| CAC FW | 55.5 | 173 | CCTCCGTTGTGATGTAACTGG |
| CAC RV | 53.5 | | ATTGGTGGAAAGTAACATCATCG |
| ACT2 Fw | 56.5 | 176 | CATTGTGCTCAGTGGTGGTTC |
| ACT2 Rv | 56.5 | | TCTGCTGGAAGGTGCTAAGTG |

**Supplemental Table 2. Structural features identified in tomato GH17 sequences.** The output of prediction platforms indicating the presence of signal peptide (SP), GPI anchor and X8 domain in the tomato sequences is summarized. Six different platforms were used for SP prediction, three platforms for GPI and three for X8 domain as described in Material and Methods. The number of glutamate (E) residues in the catalytic domain, the phylogenetic cluster the gene belongs, the type of protein according to structural predictions and the expression in fruits is also indicated.

| Gene ID | Signal peptide[1] | GPI anchor[1] | X8 domain[1] | Conserved E residues[2] | Cluster[3] | Type[4] | Expression in fruit[5] (SGN/Genevestigator) |
|---|---|---|---|---|---|---|---|
| **Solyc04g011730** | 5 | 0 | 3 | 0 | β | 3 | |
| **Solyc01g060010** | 0 | 0 | 0 | 1 | γ | 0 | |
| **Solyc11g065300** | 0 | 0 | 0 | 1 | γ | 0 | |
| **Solyc01g008610** | 3 | 0 | 0 | 1 | γ | 1 | |
| **Solyc11g065290** | 5 | 0 | 0 | 1 | γ | 1 | |
| **Solyc04g011720** | 5 | 0 | 3 | 1 | β | 3 | |
| **Solyc05g015160** | 5 | 0 | 3 | 1 | β | 3 | |
| **Solyc11g072230** | 6 | 2 | 0 | 1 | α | 4 | |
| **Solyc01g005830** | 2 | 2 | 3 | 1 | α | 5 | yes |
| **Solyc02g080660** | 5 | 2 | 3 | 1 | α | 5 | yes |
| **Solyc04g007910** | 6 | 2 | 3 | 1 | α | 5 | |
| **Solyc05g015170** | 3 | 1 | 3 | 1 | β | 5 | |
| **Solyc08g005000** | 5 | 3 | 3 | 1 | α | 5 | yes |
| **Solyc12g008580** | 6 | 2 | 3 | 1 | α | 5 | |
| **Solyc11g065280** | 0 | 0 | 0 | 2 | γ | 0 | |
| **Solyc01g008620** | 5 | 0 | 0 | 2 | γ | 1 | |
| **Solyc01g059980** | 5 | 0 | 0 | 2 | γ | 1 | |
| **Solyc02g086700** | 4 | 0 | 0 | 2 | γ | 1 | |
| **Solyc03g025650** | 5 | 0 | 0 | 2 | γ | 1 | |
| **Solyc04g016470** | 3 | 0 | 0 | 2 | γ | 1 | yes |
| **Solyc10g079860** | 5 | 0 | 0 | 2 | γ | 1 | |
| **Solyc00g202560** | 0 | 0 | 2 | 2 | γ | 2 | |
| **Solyc06g073710** | 0 | 1 | 0 | 2 | α | 2 | |
| **Solyc01g109570** | 6 | 0 | 3 | 2 | α | 3 | yes |
| **Solyc06g076170** | 5 | 0 | 3 | 2 | α | 3 | |
| **Solyc07g005330** | 5 | 0 | 3 | 2 | α | 3 | yes |
| **Solyc07g049370** | 5 | 0 | 3 | 2 | α | 3 | yes |
| **Solyc09g057630** | 5 | 0 | 3 | 2 | α | 3 | yes |
| **Solyc12g014420** | 5 | 0 | 3 | 2 | α | 3 | |
| **Solyc12g040860** | 5 | 0 | 3 | 2 | α | 3 | |
| **Solyc12g098560** | 5 | 0 | 3 | 2 | α | 3 | |
| **Solyc02g070450** | 5 | 2 | 0 | 2 | α | 4 | |
| **Solyc03g082900** | 5 | 3 | 0 | 2 | α | 4 | |
| **Solyc04g080260** | 6 | 1 | 0 | 2 | α | 4 | yes |
| **Solyc07g008150** | 5 | 2 | 0 | 2 | α | 4 | yes |
| **Solyc08g083310** | 6 | 1 | 0 | 2 | α | 4 | yes |
| **Solyc11g068440** | 6 | 3 | 0 | 2 | α | 4 | yes |
| **Solyc11g071520** | 6 | 1 | 0 | 2 | α | 4 | |
| **Solyc12g055840** | 6 | 2 | 0 | 2 | α | 4 | yes |
| **Solyc03g058450** | 6 | 3 | 3 | 2 | β | 5 | |
| **Solyc04g015190** | 3 | 2 | 3 | 2 | β | 5 | yes |
| **Solyc04g051590** | 6 | 2 | 3 | 2 | β | 5 | |

| | | | | | | | |
|---|---|---|---|---|---|---|---|
| **Solyc05g006210** | 4 | 2 | 3 | 2 | β | 5 | |
| **Solyc05g025500** | 6 | 3 | 3 | 2 | β | 5 | yes |
| **Solyc05g054440** | 1 | 1 | 3 | 2 | α | 5 | |
| **Solyc07g017730** | 2 | 2 | 3 | 2 | β | 5 | |
| **Solyc08g074390** | 4 | 2 | 3 | 2 | β | 5 | yes |
| **Solyc10g078510** | 5 | 2 | 3 | 2 | β | 5 | |
| **Solyc11g012030** | 1 | 1 | 3 | 2 | β | 5 | |
| **Solyc12g019890** | 6 | 2 | 3 | 2 | β | 5 | yes |

[1]The numbers indicate the number of platforms predicting the presence of these features.
[2]The number of glutamate (E) residues identified in the hydrolytic catalytic domain.
[3]The cluster in the phylogenetic tree where the gene is grouped is indicated to figure 1.
[4]The following classification is used: type 0 = no predicted SP, X8 or GPI-anchor domains; type 1 = only SP predicted; type 2 = only predicted X8 and/or GPI-anchor domains; type 3 = predicted SP and X8; type 4 = predicted SP and GPI; type 5 = predicted SP, GPI-anchor and X8.
[5]Yes indicates genes which are significantly expressed in pericarp tomato fruit according to SGN and Genevestigator.

**Supplementary Table 3. Absolute values of expression for GH17 genes expressed in tomato pericarp from anthesis to red ripe**. Data is extracted from the SGN-TEA database. DPA = days post-anthesis.

| GENE NAME | Anthesis | 5 DPA | 10 DPA | 20 DPA | 30 DPA | Mature green stem | Mature green equatorial | Mature green stylar | Breaker stem | Breaker equatorial | Breaker stylar | Pink stem | Pink equatorial | Pink stylar | Light red | Red ripe |
|---|---|---|---|---|---|---|---|---|---|---|---|---|---|---|---|---|
| **Solyc04g015190** | 15.5 | 19.54 | 20.9 | 19.26 | 32.76 | 27.27 | 31.96 | 33.16 | 22.46 | 25.97 | 25.4 | 16.34 | 12.71 | 13.13 | 10.39 | 12.54 |
| **Solyc04g016470** | 0 | 0.04 | 0.05 | 16.72 | 66.42 | 76.95 | 96.15 | 89.32 | 409.93 | 366.42 | 321.25 | 396.6 | 363.75 | 400.09 | 315.35 | 99.37 |
| **Solyc04g080260** | 19.46 | 15.78 | 25.67 | 36.78 | 49.6 | 40.67 | 49.85 | 48.32 | 37.23 | 43.2 | 48.79 | 30.77 | 34.91 | 29.2 | 27.84 | 29.42 |
| **Solyc11g068440** | 2.7 | 18.3 | 26.26 | 38.33 | 63.27 | 161.96 | 93.47 | 74.91 | 151.6 | 78.44 | 66.72 | 101.56 | 78.25 | 68.44 | 75.71 | 62.12 |
| **Solyc01g109570** | 1.8 | 45.92 | 70.27 | 50.7 | 57.41 | 120.1 | 80.77 | 61.64 | 83.95 | 53.39 | 54.6 | 48.53 | 40.69 | 33.47 | 30.59 | 40.81 |
| **Solyc07g008150** | 10.19 | 11.09 | 12.98 | 17.36 | 23.73 | 29.64 | 27.3 | 27.61 | 23.29 | 22.46 | 20.58 | 13.68 | 11.36 | 10.37 | 9.71 | 8.44 |
| **Solyc05g025500** | 7.71 | 34.48 | 29.96 | 16.73 | 16.74 | 18.73 | 19.23 | 18.89 | 15.29 | 15.75 | 16.49 | 8.22 | 7.98 | 8.34 | 7.14 | 6.1 |
| **Solyc01g005830** | 17 | 72.28 | 64.1 | 31.03 | 25.47 | 23.84 | 27.69 | 26.76 | 19.45 | 19.79 | 21.32 | 11.39 | 12.17 | 10.81 | 10.77 | 8.86 |
| **Solyc02g080660** | 16.07 | 166.36 | 98.57 | 31.22 | 22.63 | 22.77 | 22.83 | 22.7 | 15.76 | 15.83 | 14.22 | 11.79 | 9.93 | 10.75 | 9.94 | 9.29 |
| **Solyc08g005000** | 11.9 | 77.55 | 73.43 | 23.79 | 15.4 | 7.52 | 6.87 | 6.88 | 3.2 | 2.94 | 3.5 | 1.33 | 1.91 | 1.47 | 1.41 | 2.38 |
| **Solyc07g005330** | 33.9 | 333.07 | 325.6 | 106.72 | 82.36 | 155.78 | 100.23 | 81.72 | 81.07 | 48.32 | 46.07 | 41 | 36.22 | 27.27 | 31.79 | 30.65 |
| **Solyc07g049370** | 51.63 | 240.58 | 120.28 | 32.22 | 13.2 | 10.37 | 8.3 | 8.14 | 1.86 | 1.6 | 1.91 | 0.18 | 0.24 | 0.26 | 0.03 | 0.04 |
| **Solyc08g083310** | 18.6 | 46.49 | 64.07 | 40.12 | 26.23 | 34.63 | 29.76 | 26.65 | 7.88 | 6.76 | 7.56 | 1.72 | 2.14 | 1.28 | 1.49 | 1.32 |
| **Solyc12g055840** | 805.68 | 226.86 | 281.55 | 196.31 | 128.18 | 120.26 | 130.33 | 124.4 | 48.65 | 60.16 | 56.08 | 4.9 | 4.08 | 3.73 | 1.05 | 0.45 |
| **Solyc08g074390** | 23.56 | 4.06 | 11.43 | 16.73 | 18.92 | 19.87 | 22.45 | 21.06 | 14.99 | 14.81 | 15.24 | 9.05 | 9.01 | 5.19 | 5.33 | 6.38 |
| **Solyc12g019890** | 3.66 | 11.33 | 9.2 | 6.72 | 8.87 | 17.58 | 12.81 | 9.69 | 18.13 | 10.2 | 7.35 | 12.33 | 8.44 | 4.96 | 3.93 | 3.76 |
| **Solyc09g057630** | 6.53 | 15.77 | 15.14 | 15.19 | 13.86 | 18.02 | 17.4 | 14.34 | 9.53 | 9.58 | 7.92 | 3.73 | 3.19 | 2.29 | 2.04 | 1.36 |
